# Targets of influenza Human T cell response are mostly conserved in H5N1

**DOI:** 10.1101/2024.09.09.612060

**Authors:** John Sidney, A-Reum Kim, Rory D. de Vries, Bjoern Peters, Philip S. Meade, Florian Krammer, Alba Grifoni, Alessandro Sette

## Abstract

Frequent recent spillovers of subtype H5N1 clade 2.3.4.4b highly pathogenic avian influenza (HPAI) virus into poultry and mammals, especially dairy cattle, including several human cases, increased concerns over a possible future pandemic. Here, we performed an analysis of epitope data curated in the Immune Epitope Database (IEDB). We found that the patterns of immunodominance of seasonal influenza viruses circulating in humans and H5N1 are similar. We further conclude that a significant fraction of the T cell epitopes is conserved at a level associated with cross-reactivity between avian and seasonal sequences, and we further experimentally demonstrate extensive cross-reactivity in the most dominant T cell epitopes curated in the IEDB. Based on these observations, and the overall similarity of the neuraminidase (NA) N1 subtype encoded in both HPAI and seasonal H1N1 influenza virus as well as cross-reactive group 1 HA stalk-reactive antibodies, we expect that a degree of pre-existing immunity is present in the general human population that could blunt the severity of human H5N1 infections.

## Introduction

Influenza viruses (influenza A viruses H1N1 and H3N2, and influenza B viruses) cause seasonal infections in humans, leading up to 650,000 deaths per year globally. In addition, influenza A viruses (IAV) cause pandemics in irregular intervals that can cause millions of deaths. The most prominent example is the H1N1 pandemic of 1918, which caused an estimated 20 to 100 million deaths. Since then, influenza pandemics have occurred in 1957 (H2N2), in 1968 (H3N2) and in 2009 (H1N1). Typically, these pandemics were caused by new reassorting viruses that contain genomic segments of human or mammalian influenza viruses, as well as segments from avian influenza viruses coding for surface glycoproteins to which humans were naive (a process known as ‘antigenic shift’). IAV are classified in subtypes according to antigenic and phylogenetic relationships of the hemagglutinin (HA) and neuraminidase (NA) antigens. The highly pathogenic avian influenza (HPAI) virus of the H5N1 subtype is presently amongst the top viruses of pandemic concern according to the WHO and the NIAID pandemic preparedness plan^1,2^. While the ‘HPAI’ phenotype refers to pathogenicity in poultry, the H5N1 subtype has also been associated with severe disease in humans following animal-to-human transmission, even though human-to-human transmission has thus far been exceedingly rare^3-5^.

HPAI H5N1 possesses a polybasic cleavage site in its HA that can be cleaved by furin-like proteases, which allows replication outside of the respiratory tract. This trait is a major virulence factor in avian species as well as mammals. Infections with H5N1 in humans are usually severe, with a case fatality rate around of 50%, although asymptomatic infections may be underestimated^6^. There is a concern that the human population does not have a high degree of pre-existing immunity to HPAI of the H5N1 subtype, contributing to a susceptibility to severe disease. Indeed, cross-reactivity of hemagglutination inhibiting antibodies (which are the main correlate of protection for influenza viruses^7^) with H5 is hardly induced by infections with seasonal IAV^8,9^. However, the N1 component is shared between seasonal H1N1 and H5N1. Pre-existing immunity to N1 is therefore expected to be high^10,11^, and antibodies to NA are a correlate of protection^12^. Furthermore, it can be assumed that a fraction of the population has anti-HA stalk antibodies that cross-react between group 1 HAs like H1 and H5 and may be beneficial^13,14^. There is also an expectation that some of the T-cell epitopes in NP, M1 and other proteins are highly conserved, and there is already some evidence of cross-reactive T cells in the human population^15-18^.

Previous exposure by infection and/or vaccination to a given IAV subtype or clade influences immune responses to a different subtype or clade. This phenomenon is referred to as original antigenic sin^19,20^ and/or antigenic imprinting, antigenic seniority or back-boosting^21^. Whether this antigenic imprint is beneficial or detrimental might depend on the specific context, subtypes and type of adaptive immune response considered. Pre-existing immune memory to seasonal (s)H1N1 might have been a contributing factor in mitigating disease severity in the context of infection with the 2009 swine-origin pandemic (p)H1N1. Indeed, pH1N1 neutralizing antibodies were found in individuals born before 1957 presumably reflecting exposure to H1N1 circulating in humans before that date, and those age brackets were associated with lower disease severity^22,23^. Similarly, it was found that individuals first exposed to group 1 viruses like H1N1 and H2N2 were later in life protected from severe outcomes with H5N1 (also group 1 HA). Vice versa, initial infection with group 2 viruses like H3N2 viruses later protected from H7N9 (also group 2 HA). It has been hypothesized that this effect may be due to HA stalk-reactive antibodies^24^. Additional data indicated a beneficial effect of cross-reactive T cell immunity, as pre-existing T cells in the general population as a function of age was invoked as an explanation of differential disease severity during the 2009 H1N1 pandemic. A similar beneficial effect was observed in controlled human influenza challenge models^25-29^. Another example of beneficial cross-reactive immunity is the considerable resistance to H3N2 in 1968, observed in individuals with high N2-specific antibody levels from previous H2N2 infections^30,31^.

The Influenza virus has a negative-sense segmented RNA genome that encodes at least eleven different proteins, with HA and NA being exposed on the surface of the virus, and therefore readily accessible for antibody recognition^32^. The matrix (M1 and M2) proteins are also relatively abundant in viral particles or infected cells, and M2 is partially accessible to antibodies (M2e). The internal proteins found in association with the ribonucleoprotein (RNP) complex are nucleoprotein (NP), and the polymerase subunits (PB2, PB1 and PA). Additional proteins expressed by IAV are the non-structural proteins 1 (NS1) and the nuclear export protein (NEP). Different accessory proteins like PA-X and PB1-F2 are expressed in infected cells as well^33,34^. Internal influenza virus proteins can be targeted by antibodies but these antibodies are largely inconsequential in terms of preventing infection and disease^21,35^. However, these internal proteins are recognized by CD4 and CD8 T cells, which is consistent with observations from other viral families, and reflective of basic differences in antigen recognition between humoral and cellular adaptive immunity. Of relevance, in deciphering the potential role of cross-reactive immunity in the case of HPAI in general and H5N1 in particular, it has been noted that the internal proteins are fairly well conserved.

## Results

### Immunodominance hierarchy of H5N1 epitopes

An analysis of human humoral and cellular response data curated in the Immune Epitope Database (IEDB^36^) was performed, specifically for epitopes in HPAI of the H5N1 subtype. This first analysis focused on antigens from all H5N1 viruses, as the IEDB curates published reports from the scientific literature and no information is as yet available related to defined epitopes from the specific clades of current concern, such as the 2.3.4.4b clade. We further analyzed data from other IAV subtypes to determine whether the same regions are recognized in different subtypes, and understand the degree of conservation of epitope regions observed, to project the potential for cross-reactivity. This analysis is further complicated by the fact that HPAI H5N1 has reassorted extensively and has acquired a diversity of genomic segments encoding for internal proteins from many other avian influenza viruses, leading to a large number of different genotypes^37^.

As a first step, we queried the IEDB to retrieve H5N1 epitope data by specifying as organism the H5N1 subtype (H5N1) (organism ID:102793, H5N1), and selecting positive assays only. Three independent queries were performed to separately retrieve B cell (B cell assay data only), CD4 T cell (MHC restriction type: class II) or CD8 T cell (MHC restriction type: class I) restricted epitopes. Combined, these three queries retrieved a total of 486 epitopes derived from over 2,000 different assays reported in 139 peer-reviewed reports and submissions (as of May 2024). Of those, 298 were associated with responses in a human host, 190 in a murine host, and 15 in other species, including dogs and chickens. (The number of epitopes does not reflect the sum of the epitopes, as some epitopes have been identified in multiple species.) Out of these, 74 epitopes were associated with CD8 T cell responses, 202 with CD4 T cell responses and 210 with B cell/antibody responses.

We next investigated the immunodominance hierarchy of B cell, and CD4 and CD8 T cell responses as a function of the viral antigens. **Fig. 1a** summarizes the three types of responses as a function of each influenza virus antigen and lists the relative percentage of epitopes to address the most immunodominant antigens recognized. A limitation of this approach lies in the fact that it is based on reported positive data, where most of the literature to date has tended to focus only on selected antigens. Thus, it is not possible to exclude that in some cases a low reported number of epitopes might reflect the fact that responses to certain antigens have not been thoroughly characterized.

**Fig. 1.**
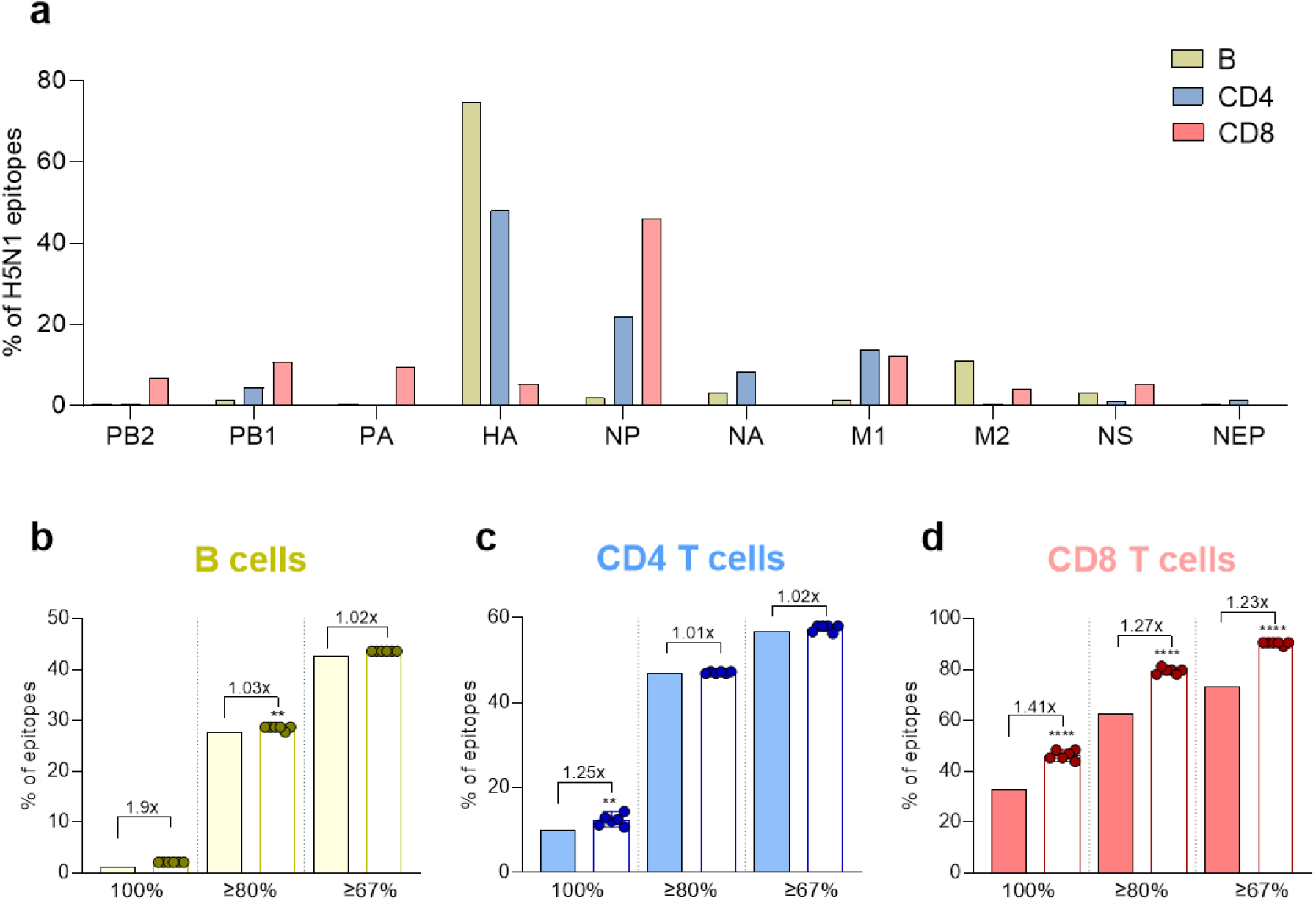
**a**, Percentage of H5N1 B cell, and CD4 and CD8 T cell epitopes per influenza virus antigen. No epitopes were reported for PA-X and other accessory proteins. **b-d**, Conservation of H1N1, H2N2 and H3N2 epitopes in avian isolates. The percentages of epitopes are shown based on the level of conservation greater or equal to 100%, 80%, and 67%. B cells (yellow, **b**). CD4 T cells (blue, **c**). CD8 T cells (red, **d**). Conservation with prototype H5N1 sequence (solid bars) is compared with the distribution of conservation for six recent isolates by one sample T-test. Fold change is calculated as the ratio of the average of the novel isolates Vs the prototype one. **p<0.01; ****P<0.0001.

In terms of B cell responses, the majority of the 210 epitopes were derived from HA (74.8%; 70.5% as a monomer and 4.3% as tertiary/quaternary epitopes on the trimer), followed by M2 (11%), NA and NS1 (3.3% each). The NP, PB1-F2, PB1, M1, PB2, NEP, and PA antigens made up for the remaining 10.9% of B cell epitopes. In terms of CD4 T cell/class II-restricted responses, the majority of the 202 epitopes were derived from HA (48%), followed by NP (21.8%), M1 (13.9%), and NA (8.4%). The PB1, M2, NS1, NEP, and PB2 antigens made up for the remaining 8.5% of the CD4 T cell epitopes. For CD8 T cell/class I-restricted responses, the majority of the 74 epitopes were derived from NP (45.9%), followed by M1 (12.2%), PB1 (10.8%), PA (9.5%), and PB2 (6.8%). The HA, NS1, and M2 antigens made up for the remaining 14.9% of the CD8 T cell epitopes. In conclusion, different patterns of immunodominance as a function of adaptive responses were observed. More specifically, the surface-exposed HA antigen was dominant in terms of B and CD4 T cell targets, even though this should be interpreted with caution because research is biased towards mostly focusing on HA. In general, very few NA mAbs have been characterized. Furthermore, the internal NP antigen was a dominant target of CD8 T cell responses.

### Immunodominance pattern between H5N1 and H1N1, H3N2, and H2N2

Estimating the potential impact of a widespread H5N1 outbreak requires understanding the extent to which pre-existing immunity originating from prior vaccinations and infections cross-reacts to H5N1. Accordingly, we surveyed the IEDB and extracted B cell, and CD4 and CD8 T cell epitope data pertaining to the influenza H1N1, including pre-pandemic seasonal H1N1 and 2009 pandemic H1N1 (Organism ID:114727), H3N2 (Organism ID:119210) and H2N2 (Organism ID:114729) subtypes, which have historically circulated in humans^38,39^, and for which immune memory is therefore expected to be prevalent. As expected, more data was available compared to H5N1, which allowed us to restrict the query to human epitope data and still obtain enough data for meaningful analysis. We understand that significant differences exist in terms of pre- and post-2009 pandemic H1N1 sequences; however, the IEDB captures data ranging from 1983 to date, and even limiting the data by year of publication it would still be unclear for each reported data when each exposure occurred and associated with which sequence.

Using the assay search panel to filter by the type of experiment performed returned a total of 2,245 human epitopes. Of these, 973 epitopes were associated with T cell assays (171 CD8 and 676 CD4 and the remaining non-assigned) and 366 with B cell/antibody assays; 1,112 epitopes have been characterized for MHC binding capacity on the basis of the use of purified MHC (662), peptide elution assays (498), or X-ray structures (29). As mentioned above, the number of total epitopes does not reflect the sum of the epitopes in each category, because some epitopes were associated with multiple categories.

The data was inspected for patterns of B cell, and CD4 and CD8 T cell immunodominance as a function of viral antigen and compared to the H5N1 data shown above. We plotted the % of epitopes derived from each antigen and recognized in H5N1 versus H1N1, H3N2, and H2N2 influenza virus for CD4 and CD8 T cell responses **(Fig. 2)**.

**Fig. 2.**
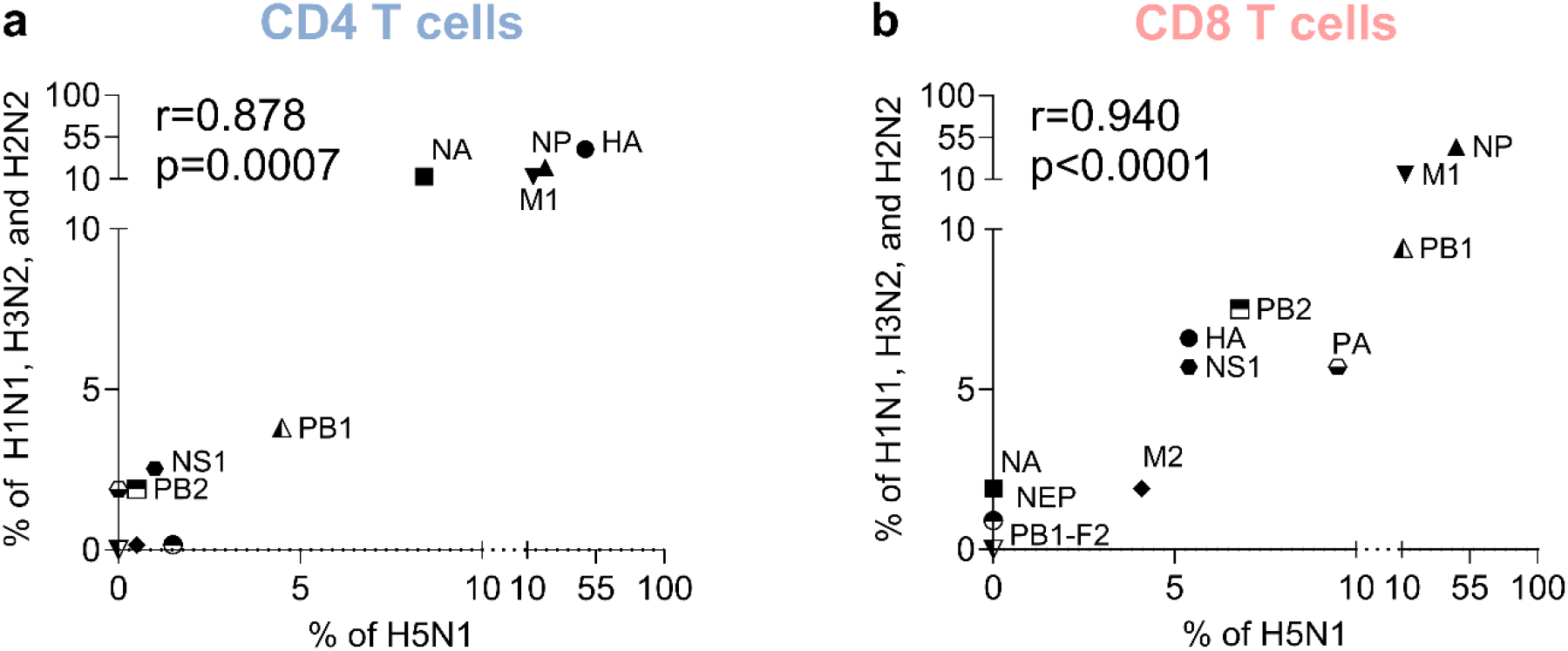
Antigen immunodominance comparison between avian (H5N1) and non-avian (influenza H1N1, H3N2 and H2N2) subtypes for CD4 (**a**) and CD8 T (**b**) cells. Spearman correlations were calculated, and two-tailed p-values shown in each panel.

T cell responses targeted a varied number of antigens, thus allowing to test for similarity in the pattern of antigenic immunodominance at the T cell level. For both CD4 (*R=0*.*878, p=0*.*0007)*, and CD8 (*R=0*.*940, p<0*.*0001)* T cells, the results revealed a high degree of correlation in immunodominance patterns between HPAI H5N1 and seasonal IAV (**Fig. 2**). In the context of B cell responses, in both cases most of the data is associated with HA, and the number of reported epitopes for other antigens is minimal (**Fig. 1a**), rendering a correlation analysis statistically unmeaningful. Nevertheless, antibody responses in HPAI H5N1 and seasonal IAV are similar in terms of predominantly targeting the HA antigen, even though this is likely to at least in part reflect research bias. The relative scarcity of defined epitopes derived from NA curated in the IEDB was surprising, given the fact this antigen is clearly associated with meaningful protective responses^12^. Overall, this indicates a fundamental similarity between the antigen specificity of T and B cell immune responses across IAV strains.

### H1N1, H3N2, and H2N2 epitope conservation across recent H5N1 isolates

Several studies in different virus families, including coronaviruses and flaviviruses, demonstrated that cross-reactivity at the T cell level is predictable based on sequence similarity^40-43^. A threshold of 67% homology was experimentally demonstrated as being associated with cross-reactivity for CD4 T cells^40,41,43^, and a threshold of 80% homology was experimentally demonstrated as being associated with cross-reactivity for CD8 T cells^42^. In contrast, cross-reactivity at the antibody level between different viral antigens is in general sensitive to both sequence variation and conformational changes, and therefore is in general more difficult to predict.

To study the potential for cross-reactivity between seasonal IAV- and HPAI H5N1-specific immune responses, we queried the IEDB for the most dominant (frequently recognized) human antibody, and CD4 and CD8 T cell epitopes derived from H1N1, H2N2 and H3N2 subtypes. Only linear epitopes were considered, since no tool is currently available to easily allow for conservation analysis of discontinuous epitopes, and our search was limited to identifying only epitopes associated with positive responses in humans. CD4 T cell, CD8 T cell, and antibody epitopes were analyzed separately. In all, this corresponded to a total of 33 possible response type/antigen combinations (3 types of epitopes X 11 antigens). For each combination, we used the ImmunomeBrowser to download the associated epitope data. We further filtered for canonical size in the case of T cell epitopes, considering 8-11 residues for CD8, and 12-25 for CD4; no size filter was applied for antibody epitopes. Epitopes with reported positive responses in at least 2 subjects and associated with an overall Response Frequency (RF) value of 0.1 or greater^44^ were defined as dominant. The results, summarized by antigen in **Supplementary Table 1**, identified a list of 224 CD4 epitopes, 64 CD8 epitopes, and 94 B cell epitopes.

Next, utilizing the sequence conservation tool provided in the IEDB Analysis Resource (IEDB-AR^45^), we examined the degree of sequence conservation of these human seasonal IAV dominant epitopes in the HPAI H5N1 isolate A/Anhui/1/2005 H5N1, taken as a prototype HPAI of the H5N1 subtype genome. The results (**Table 1**) indicate that 9.8% of CD4 and 32.8% CD8 epitopes were fully conserved (100% sequence identity). By contrast, only one (1.1%) of the B cell epitopes was fully conserved. Furthermore, it was noted in total 56.7% of CD4 and 62.5% CD8 T cell epitopes are conserved above the 67% and 80% sequence identity thresholds associated with experimentally verified cross-reactivity for CD4 and CD8 T cell responses, respectively^40-43^. We also analyzed conservation in 6 different recent clade 2.3.4.4b H5N1 isolates from the Americas from several animal species during the 2023-2024 period and noted an overall similar level of conservation (**Table 1**). These virus isolates represent 5 genotypes and include the B3.13 genotype that currently circulating in dairy cattle and causing concern over its pandemic potential.

**Table 1.** Conservation of H1N1, H2N2 and H3N2 epitopes in different H5N1 sequences. The number and percentage of epitopes are shown based on the level of conservation equal to 100%, or greater than 80% and 67%.

| Conservation | Prototype<br>(A/Anhui/1/2005) | A/chicken/<br>New<br>Jersey/23-027703-001-original/20<br>23 (B3.3) | A/chicken/M<br>aine/24-007946-001/2024<br>(C2.1) | A/snow<br>goose/Cali<br>fornia/24_0<br>04881-004/2024<br>(B3.2) | A/South<br>American<br>sea<br>lion/Argenti<br>na/RN-PB011/2023<br>(B3.2) | A/black<br>vulture/Pen<br>nsylvania/2<br>3-015076-001-original/202<br>3 (B1.1) | A/dairy<br>cow/Texas/<br>24_008749-006-original/202<br>4 (B3.13) |
| --- | --- | --- | --- | --- | --- | --- | --- |
| **CD4 (n=224)** |  |  |  |  |  |  |  |
| 100% | 22(9.8%) | 27(12.1%) | 25(11.2%) | 24(10.7%) | 28(12.5%) | 32(14.3%) | 29(12.9%) |
| 80% | 105(46.9%) | 106(47.3%) | 105(46.9%) | 105(46.9%) | 106(47.3%) | 105(46.9%) | 106(47.3%) |
| 67% | 127(56.7%) | 130(58%) | 130(58%) | 130(58%) | 130(58%) | 126(56.3%) | 127(56.7%) |
| **CD8 (n=64)** |  |  |  |  |  |  |  |
| 100% | 21(32.8%) | 28(43.8%) | 31(48.4%) | 31(48.4%) | 30(46.9%) | 29(45.3%) | 29(45.3%) |
| 80% | 40(62.5%) | 50(78.1%) | 51(79.7%) | 51(79.7%) | 51(79.7%) | 52(81.3%) | 50(78.1%) |
| 67% | 47(73.4%) | 57(89.1%) | 58(90.6%) | 58(90.6%) | 58(90.6%) | 58(90.6%) | 58(90.6%) |
| **B (n=94)** |  |  |  |  |  |  |  |
| 100% | 1(1.1%) | 2(2.1%) | 2(2.1%) | 2(2.1%) | 2(2.1%) | 2(2.1%) | 2(2.1%) |
| 80% | 26(27.7%) | 27(28.7%) | 27(28.7%) | 27(28.7%) | 27(28.7%) | 26(27.7%) | 27(28.7%) |
| 67% | 40(42.6%) | 41(43.6%) | 41(43.6%) | 41(43.6%) | 41(43.6%) | 41(43.6%) | 41(43.6%) |

We next determined if the level of conservation between the prototype strain (from 2005) and across the more recent clade 2.3.4.4b H5N1 isolates/genotypes was different and decreased, which would raise concerns on the ability of the immune system primed with circulating IAV to cross-recognize the avian ones if exposed. Notably, the average conservation was similar or even significantly increased as compared to the prototype (**Fig. 1b-d**).

In general, it is appreciated that the targets of humoral and cellular immune responses only partially overlap. In the case of influenza virus, antibody responses target the more variable and surface-accessible HA and NA antigens. In the case of clade 2.3.4.4b H5N1, based on the sequence similarity of the N1 component, extensive cross-reactivity at the level of antibody response is expected. There is data to support this notion^10,11^. The IEDB only reports defined epitopes and there are reports describing at least four different supersites (HA central stalk, HA anchor, active site of NA, sialic acid binding site of HA) that are targeted by HA and NA cross-reactive antibodies. Furthermore, it is also expected that anti-HA stalk antibodies induced by H1N1 will cross-react to clade 2.3.4.4b H5 HA since they were detectible in humans against clade 2.3.4.4 H5 HA dependent on age and exposure history^13^. However, the globular head domain of HA (which is the main target of neutralizing antibodies) is very plastic and the H5 head domain differs drastically from H1 or H3 head domains. By contrast, a significant fraction of T cell responses target virion and non-structural antigens, which are significantly more conserved, thus contributing to a higher potential for cross-reactivity of T cell responses. These mechanisms are also exemplified by the SARS-CoV-2 pandemic; sequence variation often led to escape from antibodies, but this was not observed in the context of T cell recognition^46-53^.

### T cell immune responses against the human seasonal IAV dominant epitopes and their HPAI H5N1 counterparts

To further evaluate the biological significance of the conservation of epitopes from H1N1, H2N2, and H3N2 (non-avian human strains) we synthesized the dominant T cell epitopes identified in humans **Supplementary Table 1** and their corresponding HPAI H5N1 isolate A/Anhui/1/2005 H5N1 peptides. These peptides were then pooled in four different MegaPools (MPs)^54^ corresponding to human and avian CD4 or CD8 MPs. These newly designed pools were used to measure CD4+ and CD8+ T cell responses using PBMCs from 20 healthy controls (HC) collected from Oct 2021 to July 2022. The results in **Fig 3** show significant reactivity against the human non-avian MPs using T cells derived from these donors. When we compared the antigen-specific responses of the human non-avian MPs to the avian MPs, in the case of CD4+AIM+, CD4+GranzymeB+ and CD4+IFN-γ+ T cell responses (**Fig. 3a-c**) no significant differences between non-avian and avian CD4 MPs were noted (CD4+AIM+, p=0.3385, ratio=0.886; CD4+GranzymeB+, p=0.4423, ratio=0.7818; CD4+IFN-γ+, p=0.9515, ratio=9463).

**Fig. 3.**
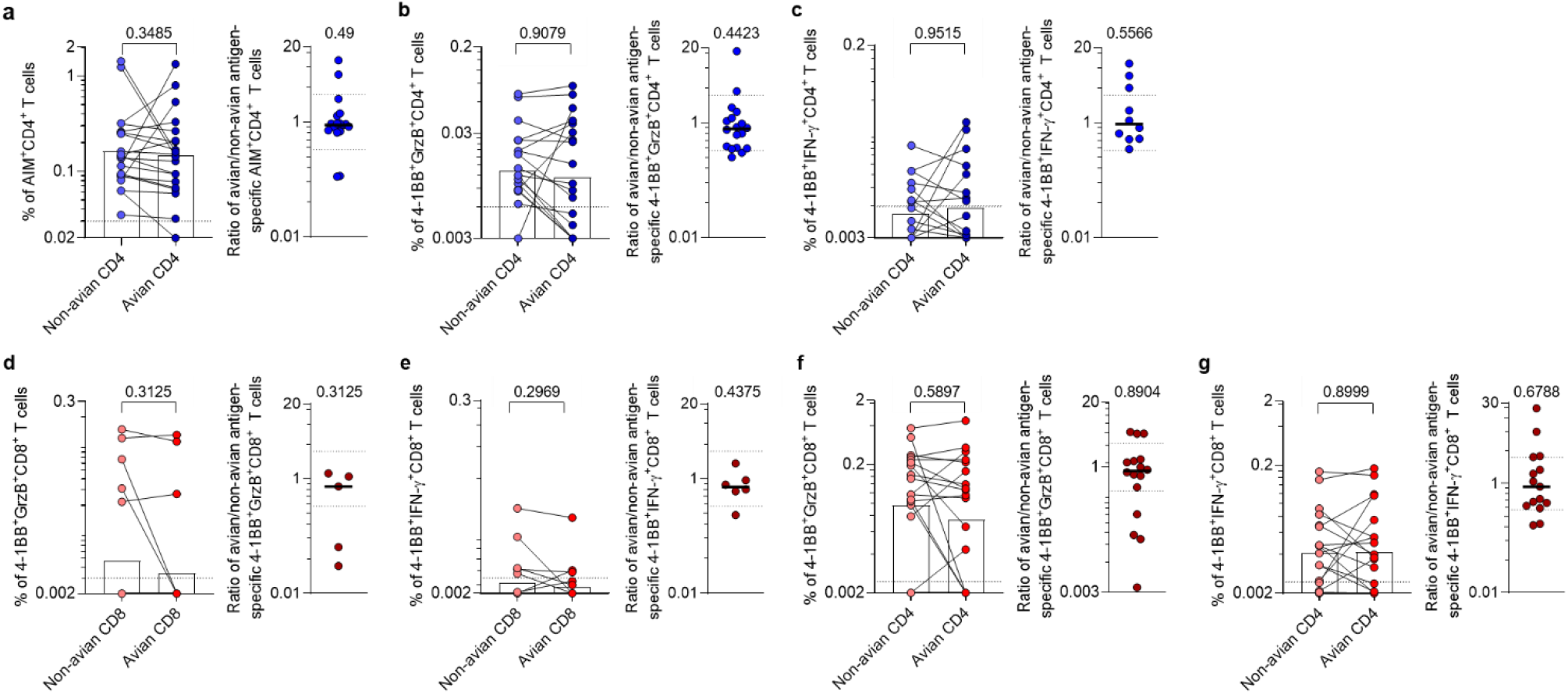
Comparison of T cell responses to non-avian (H1N1, H3N2, and H2N2) and avian (H5N1) antigens. T cell responses were assessed in PBMCs from the healthy donors. The comparison of T cell responses and the ratio between non-avian and avian MPs were shown at the left and right sides of each panel, respectively. **a**-**c**, CD4+ T cell responses to non-avian and avian CD4 MPs. CD4+AIM+ (**a**), CD4+Granzyme B+ (**b**), and CD4+IFN-γ+ (**c**) T cell responses. **d**-**e**, CD8+ T cell responses to non-avian and avian CD8 MPs. CD8+Granzyme B+ (**d**) and CD8+IFN-γ+ (**e**) T cell responses. **f**-**g**, CD8+ T cell responses to non-avian and avian CD4 MPs. CD8+Granzyme B+ (**f**) and CD8+IFN-γ+ (**g**) T cell responses. The comparisons are performed by paired Wilcoxon test and the significance of the ratio is analyzed by one-sample Wilcoxon test compared with a hypothetical median of 1. p values are listed at the top of each graph. The ratios between non-avian and avian MPs-specific responses are calculated if either value is above the LOS. Bars represent the geometric mean. The y-axis starts at the LOD, and the dotted lines indicate the LOS (bar graphs).

Because of the lower sensitivity of the AIM assays in detecting CD8+ responses, we focused on CD8+GranzymeB+ and CD8+IFN-γ+ T cell responses. No significant differences were observed between non-avian and avian CD8 MPs (CD8+GranzymeB+; p=0.3125, ratio=0.726; CD8+IFN-γ+, p=0.2969, ratio=0.7141) (**Fig. 3d-e**) despite the relatively low level of responses likely due to the limited number of epitopes in the MPs. Better overall responses were seen when the CD4 MPs, which spanned a large fraction of IAV genome and a larger number of peptides, were used to detect CD8 T cell responses (CD8+GranzymeB+; p=0.5897, ratio=0.8325; CD8+IFN-γ+, p=0.8999, ratio=0.8741) (**Fig. 3f-g**). Overall, these data demonstrate that the existing IAV-specific T cell responses are largely preserved when in terms of recognition of the corresponding HPAI H5N1 sequences.

## Discussion

A systematic analysis of human epitope conservation between HPAI H5N1 (considering the A/Anhui/1/2005 prototype) and seasonal IAV sequences revealed that 57% of CD4 and 63% of CD8 T cell epitopes, respectively, are either identical or conserved to a level of sequence identity that is expected to be conducive to T cell cross-reactivity. Overall, this analysis predicted that pre-existing T cell immune memory would, to a large extent, cross-recognize avian influenza viruses, should more widespread human infections start to occur. This prediction was experimentally verified testing responses from human T cells to non-avian IAV and their HPAI H5N1 counterparts.

Antibody-antigen interactions are in general sensitive to changes in antigenic structure and sequence as compared to T cell recognition, and conversely there are cases where the sequence is very different but antibodies are still binding and neutralizing. While no method relating sequence and structural similarity is currently available to easily predict antibody cross-reactivity, our analysis reveals that compared to T cell epitopes, fewer antibody epitopes are fully conserved in HPAI H5N1. While several studies report that specific monoclonal antibodies defined following exposure to non-H5N1 influenza viruses can cross-react with H5N1 avian influenza virus antigens^55-63^, the functional significance in terms of antibody cross-reactivity between HPAI H5N1 and seasonal IAV at the polyclonal level, or even population level, remains to be established. Cross-reactive immunity at the level of B cell responses can influence immune responses to novel strains. This may result in muting or skewing immune responses towards non-protective responses in an “original antigenic sin”-like fashion, but, conversely, could also be beneficial^21,24^.

The next logical question is whether pre-existing cross-reactive T cell immune memory would be beneficial, inconsequential or even detrimental. This issue was recently extensively debated in the case of SARS-CoV-2^64,65^, where T cell cross-recognition of different variants, despite escape from antibody recognition, was associated with prevention of severe disease and death^46-52,66^. In the case of influenza virus, there is evidence for a beneficial effect of cross-reactive T cell immunity in seasonal and pandemic H1N1, as pre-existing immunity in the general population being observed as a function of age was invoked as an explanation of differential disease severity during the 2009 pH1N1 outbreak, and in controlled human influenza virus challenge models^25-29^. In that context, it was found that 41% of the CD4 and 69% of the CD8 T cell epitopes were completely conserved between pH1N1 and previously circulating sH1N1. These levels were higher than those observed here, when studying cross-reactivity between seasonal IAV and HPAI H5N1^28^.

In the context of the SARS-CoV-2 pandemic, pre-existing immune T cell memory induced by prior infection with other coronaviruses^40^ was associated with milder disease outcome and even abortive infection^67,68^, particularly in cohorts of exposed healthcare workers^69,70^, and also associated with more vigorous responses to vaccination^71,72^. In this context, we evaluated the degree of conservation noted in panels of alpha and beta coronavirus-derived epitopes defined in pre-pandemic samples^41^ and noted that 11 out of a total of 166 (6.6%) epitopes were conserved at the 67% homology threshold level. This observation suggests that even conservation of a relatively small number of epitopes across related viruses can be of consequence in terms of modulation of disease severity.

Based on this analysis and the overall information available to date, we hypothesize that, should a widespread HPAI H5N1 outbreak in humans occur, cross-reactive T cell responses might be able to limit disease severity, but to a lower extent than what was observed in the context of 2009 pH1N1. We propose that a strategy to enhance and focus T cell immunity towards recognition of conserved and immunogenic regions within a virus family might be generally applicable to several families of potential pandemic concern^73^. We submit that this strategy might also be applicable in the case of the pandemic threat of influenza.

## Methods

### Querying influenza epitopes in IEDB database

B cell (antibody), and CD4 (Class II) and CD8 (Class I) T cell, epitopes were extracted from the IEDB database (www.IEDB.org)^36^ on May 7^th^, 2024, using the following organisms: influenza H1N1 (Organism ID:114727), H3N2 (Organism ID:119210), H2N2 (Organism ID:114729), or H5N1 (Organism ID: 102793). The query included positive assays only, with the host specified as Human. Separate queries were performed for T cell Class I and Class II epitopes by selecting either MHC restriction type Class I for CD8 epitopes or Class II for CD4 epitopes. CD8 and CD4 epitopes were further filtered on size criteria, to include only lengths of 8-11 or 12-25 residues, respectively, to comport with canonical epitopes sizes. B cell epitopes were also queried separately, with no size criteria implemented. The data reported and curated in the IEDB does not consistently identify clades, thus the granularity of the analysis was limited to the H5N1 subtype level; and the H1N1, H3N2, H2N2 subtype and human host level.

### Conservation analyses

Conservation analyses were performed using the Conservation tool available in the IEDB Analysis Resource (IEDB-AR)^74^ for non-avian epitopes based on the IEDB epitope query for H1N1, H3N2 and H2N2, as listed in Table S1. The epitopes were divided into the different influenza antigens and ran separately against the 2005 prototype HPAI H5N1 isolate EPI ISL 24603 and six different isolates collected in 2023-2024 (EPI_ISL_18731617; EPI_ISL_1909448; EPI_ISL_19094639; EPI_ISL_18945317; EPI_ISL_17964970; EPI_ISL_19151399). The resulting conservancy values were then compiled to count the number of epitopes meeting specified thresholds of conservation (100%, ≥80% and ≥67%), mirroring cutoffs previously validated experimentally to predict cross-reactivity for CD8^42,75^ and CD4^40,41,43^, respectively.

### Preparation of peptide pools

Based on conservation analyses, we synthesized sets of CD4 (n=224, non-avian CD4) and CD8 (n=64, non-avian CD8) epitopes from non-avian strains, as well as the corresponding CD4 (n=219, avian CD4) and CD8 (n=40, avian CD8) epitopes from avian strains, excluding duplicate peptides. The synthesized peptides were individually resuspended in dimethyl sulfoxide (DMSO; Sigma) at a concentration of 20 mg/ml. Subsequently, the peptides were pooled, lyophilized, and resuspended at a final concentration of 1 mg/ml in DMSO.

### Human subjects and peripheral blood mononuclear cells (PBMC) isolation

Blood samples, leftover from a previous research study and eligible to be used for the current study, were utilized from healthy adult donors (n=20) obtained from the La Jolla Institute for Immunology Clinical Core in San Diego, California (under IRB protocol VD-101). Donors were 22-61 years old (median 38), consisting of 65% females and 35% males. The cohort was 75% White/not Hispanic or Latino, 5% Asian, and 20% more than one race. The samples were collected between October 2021 and July 2022. Whole blood was drawn into heparin-coated blood bags and processed according to a previously established protocol^76^. PBMCs were isolated and cryopreserved in cell recovery media containing 10% DMSO, supplemented with 10% heat-inactivated fetal bovine serum (FBS; Hyclone Laboratories, Logan, UT). The cryopreserved PBMCs were then stored in liquid nitrogen until used in subsequent assays.

We have complied with all relevant ethical regulations pertaining to the use of human donor samples. All human subject protocols have been approved by the La Jolla Institute for Immunology’s Institutional Review Board (Federalwide Assurance Number: FWA00000032).

### Activation-induced markers (AIM) and intracellular cytokine staining (ICS) assay

The samples were tested using a combined AIM and ICS (AIM/ICS) assay, as previously described ^76^. Briefly, cryopreserved PBMCs were thawed in RPMI 1640 media supplemented with 5% human AB serum (Gemini Bioproduct) and benzonase (20 μl/10ml, EMD Milipore Corp). A total of 2×10^6^ cells per well in 96-well U-bottom plates (GenClone) were stimulated for 24h in the presence of non-avian and avian MPs (1 μg/ml). Equimolar amounts of DMSO were used in quadruplicate as a negative control, while phytohemagglutinin (PHA; 1 μg/ml, Roche) was used as a positive control. After 20h of incubation at 37°C, 5% CO2, cells were incubated in the presence of Golgi Plug (Brefeldin A) and Golgi Stop (Monensin) (both BD Bioscience) for an additional 4h to allow the accumulation of intracellular cytokines and CD137 APC (1:100; Biolegend, clone 4B4-1, Cat# 309810) to prevent re-internalization.

After 24h of cumulative stimulation, membrane staining was conducted for 30 min at 4°C using the following antibodies: CD8 BUV496 (2:100; BD Biosciences; clone RPA-T8; Cat# 612942), CD4 Brilliant Violet (BV) 605 (2:100; BD Biosciences; clone RPA-T4; Cat# 562658), and CD3 Alexa Fluor 700 (1:100; eBioscience; clone UCHT1; Cat# 56-0038-42). LIVE/DEAD Fixable NIR stain APC-ef780 (0.05: 100; Invitrogen; Cat# L34975) was used to mark viable cells. CD14 APC-ef780 (1:100; clone 61D3; eBioscience; Cat# 47-0419-42) and CD19 APC-ef780 (1:100; eBioscience; clone HIB19; Cat# 47-0199-42) were both used to exclude monocytes and B cells in the dump channel. For AIM-specific T cell identification, CD137 APC (2:100) and CD69 BV786 (1:100; BD Biosciences; clone FN50; Cat# 563834) were used for CD8+ T cells, while CD137 APC (2:100) and OX40 PE-Cy7 (2:100; Biolegend; clone Ber-ACT35; Cat# 350012) were used for CD4+ T cells.

Next, cells were fixed with 4% paraformaldehyde (PFA) solution for 10 min at 4°C, followed by permeabilization with saponin buffer and blocking with 10% human AB serum. ICS was performed for 30 min at 4°C with antibodies detecting cytokines, including IFN-γ FITC (1:100; Invitrogen; clone 4S.B3; Cat# 11-7319-82) and Granzyme B PE (1:100; Invitrogen; clone GB11; Cat# 12-8899-41). The cells were washed with PBS, resuspended in 120μL of PBS per well, and acquired by a ZE5 cell analyzer (Bio-Rad laboratories). Raw FCS files were analyzed with FlowJo software (Tree Star).

### Quantification and statistical analyses

Correlation analyses were performed by Spearman correlation; r and two-tailed p values are listed in each Figure. Conservancy analysis comparing prototype H5N1 isolate and six more recent H5N1 isolates was performed by one sample T-test. Fold-Change conservancy is calculated by dividing the average conservation for the six isolates by the prototype isolate.

The frequencies of non-avian and avian MP-specific T cells were determined using AIM and CD137+ICS gating strategies. The percentages of the quadruplicate DMSO wells were averaged, and this background was subtracted from each MP-stimulated population. The stimulation index (SI) was calculated by dividing the percentage of positive cells after MP stimulation by averaged DMSO background. For each gated population, the Limit of Detection (LOD) was determined by the two-fold lower confidence interval (CI) of the geometric mean, and the Limit of Sensitivity (LOS) was based on the median plus two-fold standard deviation (SD) of T cell reactivity in the DMSO wells. A response was considered positive if the DMSO background-subtracted value was greater than the LOS and the SI was greater than 2. Data and statistical analyses were performed in FlowJo 10 and Graphpad Prism 10.2. The comparisons are performed by paired Wilcoxon test and the significance of the ratio is analyzed by one-sample Wilcoxon test compared with a hypothetical medina of 1. The ratios between non-avian and avian MPs-specific responses are calculated if either value is above the LOS.

## Data Availability

The published article includes all data generated or analyzed during this study, and summarized in the accompanying tables, figures, and supplemental materials. Any additional information required to reanalyze the data reported in this work paper is available from the Lead Contact upon request.

## Code availability

This paper does not report the original code.

## Acknowledgments

This project has been funded in whole or in part with Federal funds from the National Institute of Allergy and Infectious Diseases, National Institutes of Health, Department of Health and Human Services, under contract no. 75N93019C00001 to A.S. and B.P. We would like to thank Randi Vita for the support on specialized IEDB queries. The Krammer laboratory receives support for work on influenza virus from the NIAID Centers of Excellence for Influenza Research and Response (CEIRR, 75N93021C00014) and NIAID Collaborative Influenza Vaccine Innovation Centers (CIVICs, 75N93019C00051). We gratefully acknowledge all data contributors, i.e., the Authors and their Originating laboratories responsible for obtaining the specimens, and their Submitting laboratories for generating the genetic sequence and metadata and sharing via the GISAID Initiative, on which this research is partially based.

## Author contributions

Conceptualization: A.G., A.S.; Methodology: J.S., A.G., B.P.; Formal analysis: J.S., R.D.dV., P.S.M, F.K., A.K., A.G.; Investigation: J.S., A.G.; Funding acquisition: A.S., B.P.; Writing: J.S., A.G., A.S, R.D.dV., B.P., F.K.

## Additional information

Supplementary information is available for this paper. Correspondence and requests for materials should be addressed to Alessandro Sette and Alba Grifoni.

